# Phinch: An interactive, exploratory data visualization framework for –Omic datasets

**DOI:** 10.1101/009944

**Authors:** Holly M. Bik, Pitch Interactive Inc.

## Abstract

Using environmental sequencing approaches, we now have the ability to deeply characterize biodiversity and biogeographic patterns in understudied, uncultured microbial taxa (investigations of bacteria, archaea, and microscopic eukaryotes using 454/Illumina sequencing platforms). However, the sheer volume of data produced from these new technologies requires fundamentally different approaches and new paradigms for effective data analysis. Scientific visualization represents an innovative method towards tackling the current bottleneck in bioinformatic workflows. In addition to giving researchers a unique approach for exploring large datasets, it stands to empower biologists with the ability to conduct powerful analyses without requiring a deep level of computational knowledge. Here we present Phinch, an interactive, browser-based visualization framework that can be used to explore and analyze biological patterns in high-throughput -Omic datasets. This project takes advantage of standard file formats from computational pipelines in order to bridge the gap between biological software (e.g. microbial ecology pipelines) and existing data visualization capabilities (harnessing the flexibility and scalability of technologies such as HTML5).

## Introduction

The advent of high-throughput sequencing data is now ushering in a veritable renaissance in biology. For the first time, we have the ability to deeply characterize the global biodiversity of historically neglected, microbial taxa via environmental sequencing approaches (investigations of bacteria, archaea, and microscopic eukaryotes using 454/Illumina sequencing platforms, e.g. Creer et al. 2010, Sogin et al 2006). However, the sheer volume of data produced from these new technologies will require fundamentally different approaches and new paradigms for effective data analysis (Akey & Shriver, 2011). Scientific visualization represents an innovative method towards tackling the current bottleneck in bioinformatics; in addition to giving researchers a unique approach for exploring large datasets, it stands to empower biologists with the ability to conduct powerful analyses without requiring a deep level of computational knowledge.

Effective, sophisticated visualization tools (taking advantage of human cognition and human information-processing capabilities) can help to link information from disparate fields, propelling scientific insight and spurring new discoveries (Munzner et al, 2006). Computer algorithms face significant difficulty in identifying simple data patterns, and thus writing algorithms for complex, subtle patterns (the type that exist in biological systems) is almost impossible. The human eye, in contrast, is very adept at spotting subtle visual patterns, able to quickly notice trends and outliers (Heer et al, 2010), especially when presented with intuitive, well-designed software tools and user interfaces (Heer & Shneiderman 2012).

The increasing scale of –Omic data (hundreds of millions of raw DNA sequences) makes it unfeasible to conduct fine-scale analyses in many existing biological software packages. In addition, the generation of high-throughput sequencing data (454, Illumina) fundamentally differs from Sanger sequencing platforms – the nature of the data requires specific computational considerations (e.g. accounting for intragenomic variation across 18S rRNA gene copies present in eukaryotic genomes, which inherently affect our interpretation of environmental sequence data; Bik et al. 2012a). Thus, given recent advances in sequencing technology, the development of exploratory data visualization approaches would be particularly well-suited for high-throughput datasets, since we do not yet understand the regularities or “classes of behavior” inherent to the underlying biological sequences (Shoresh and Wong 2012).

Here we present Phinch (http://phinch.org), an interactive, exploratory data visualization tool designed to facilitate the analysis of –Omic datasets. This software framework provides a streamlined visualization workflow and sleek user interface, aimed at enabling novel explorations of large biological datasets as part of the downstream data analysis workflow. Phinch is accessible via web browser (Google Chrome), with the software implemented locally on a user’s machine; all data processing and visual renderings are carried out using built-in javascript libraries within the browser application itself. The overarching goals of the Phinch data visualization framework have been twofold: first, to encourage researchers and public audiences to interact with and explore large –Omic datasets, regardless of computational skill level, and second, to promote effective and efficient scientific research by providing an intuitive and easy way to filter data, identify biological patterns, and export publication-quality graphics.

## Methods

Phinch is a fully open source framework written in CoffeeScript (http://coffeescript.org/), a programming language that compiles into JavaScript. The public implementation of the software is accessible at http://phinch.org, consisting of a minimalist homepage with a simple box for loading data files. The Phinch framework is released under the open source BSD-2 clause license, and the codebase is freely available on GitHub (https://github.com/PitchInteractiveInc/Phinch/). Phinch is designed to function locally on a user’s machine, within the framework of a web browser; the current software release is optimized for use in Google Chrome. The visualization framework relies on built-in javascript APIs within Chrome (e.g. IndexedDB - http://aaronpowell.github.io/db.js/) to parse files and prepare datasets for downstream visualizations. The data visualizations in Phinch are constructed using the D3.js software library (http://d3js.org/), a popular toolkit that allows for dynamic and interactive visualizations of large datasets. D3.js is a relatively new javascript library that enables rich data visualizations to be created within a web browser; this library has many features, but one strength is the ease at which scalable vector graphics (SVGs) can be created. SVG is an XML based vector image format that is supported in nearly all modern browsers. Phinch also utilizes jQuery, another javascript library that is designed to work in almost every web browser and facilitates many common web development tasks. In Phinch, jQuery is used for a wide array of features; such functions include applying various styles to different portions of the visualizations, and assigning different event handlers to facilitate executing functions when users interact with the visualizations. These two javascript libraries (D3.js and jQuery, along with a few other smaller, minor libraries) are downloaded to a user’s computer when they access the public implementation of the software at http://phinch.org. Although the software is accessible via web URL and users are promoted to load a file, for security and data privacy reasons, Phinch does not transfer user data or files over the internet by default, nor does the software index any information on remote servers.

Phinch currently supports downstream analyses of BIOM files (version 1.0 of the Biological Observation Matrix format, http://biom-format.org a JSON-formatted file type typically used to represent marker gene OTUs or metagenomic data; McDonald et al. 2012). This file type is implemented within the current stable release of QIIME (version 1.8, http://qiime.org; Caporaso et al. 2010), one of the most popular software pipelines for the processing and analysis of environmental sequence data (primarily rRNA amplicon datasets, but QIIME is also used for shotgun metagenomic data). In developing Phinch, we chose to focus on BIOM-formatted files because of the flexibility and long-term utility that this format offers for different types of biological datasets. The BIOM format is readily extensible and applicable to any type of biological matrix and associated metadata. For example, it could be used to represent OTU tables derived from environmental amplicon datasets, reported contigs from metagenomic studies with their associated ontologies, gene variant data, or even morphological character state matrices. Phinch requires only a single BIOM file as input, and from a software engineering standpoint this greatly simplifies the process of loading and parsing user datasets. We have taken advantage of built-in features offered by the BIOM file format, that allow users to embed their sample metadata and taxonomy/ontology information alongside the main biological matrix containing observation data. To utilize the Phinch framework, users must ensure that all sample metadata and associated taxonomy/ontology information is embedded within the BIOM file before loading it into the web browser. The addition of metadata and taxonomy/ontology information can be carried out with a few simple commands (Supplementary File 1 or refer to https://github.com/PitchInteractiveInc/Phinch/wiki/Quick-Start), using either the standalone BIOM format toolkit (https://pypi.python.org/pypi/biom-format/), or the BIOM commands available in QIIME (version 1.7 or higher).

Extensive and updated documentation for the Phinch visualization framework is available on the wiki page of the GitHub software repository: https://github.com/PitchInteractiveInc/Phinch/wiki

## Results

The features and visualizations within the Phinch framework were developed in order to pick up where the functionality of current bioinformatics tools ends. Phinch assumes that users have already performed upstream processing and data analysis workflows (e.g. preparing raw data and applying the necessary quality controls, filtering/clustering data, appending data points with taxonomy/ontology information), and embedded the necessary sample metadata and taxonomy/ontology information into the BIOM file (Supplementary File 1).

After loading a data file, users are initially directed to a “Parser Window” (Figure 1) that summarizes the observation data and sample metadata stored in the BIOM file. This window displays a summary of the biological matrix (sample names, number of samples, observations per sample), and allows users to sort and filter samples based on metadata values. Slider bars and checkboxes on the left-hand side of the parser window provide a quick, intuitive way to define specific criteria; samples meeting the resulting criteria are filtered in real time as numerical ranges and metadata categories are adjusted by the user.

**Figure 1:**
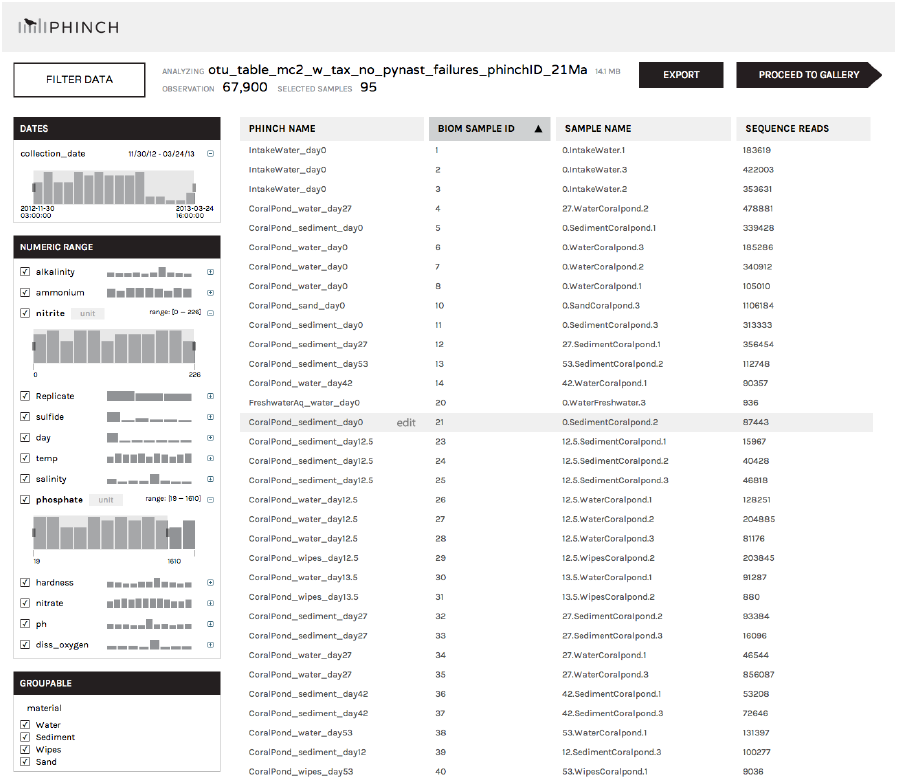
Parser Window for sample sorting and metadata filtering. After loading a file into the Phinch framework, the software imports sample IDs, observation data (e.g. sequence reads, OTUs, gene occurrence data, etc.), and associated metadata, into the “Parser Window”. Users are able to sort and filter samples in real time, using the slider bars on the left hand side of the browser window as well as the column headers. Clicking on the “Proceed to Gallery” button will load the selected samples into the downstream data visualizations.

After filtering samples using the Parser Window, users can access the data visualizations by clicking the button reading “Proceed to Gallery” (Figure 1). The Phinch framework currently offers five distinct visualizations tools (Figure 2), as follows:

**Figure 2:**
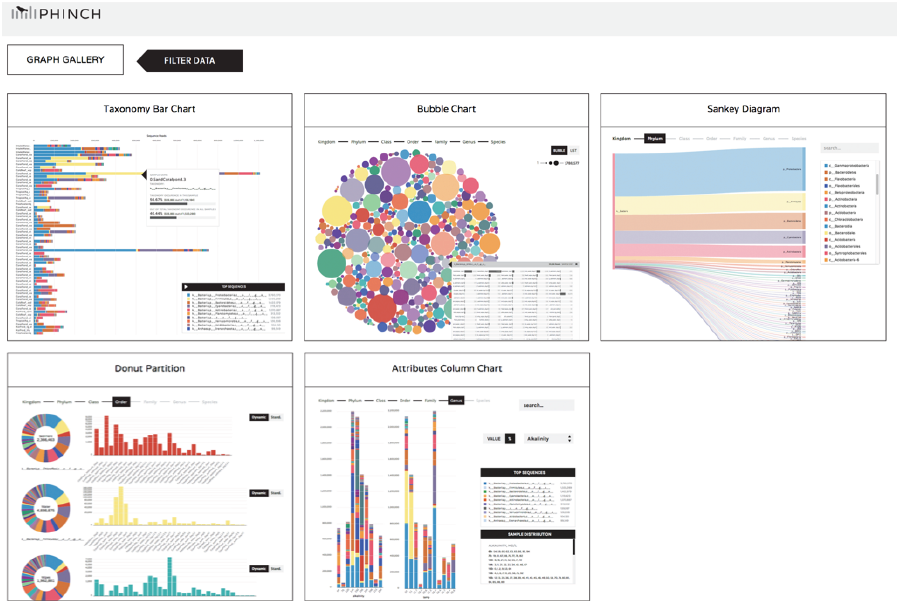
Visualization Gallery in Phinch. After manipulating data in the Parser Window, users are taken to the main visualization gallery, which offers a choice of different data visualization tools. Five interactive visualization have been implemented in the current software release: Taxonomy Bar Charts, Bubble Charts, Sankey Diagrams, Donut Partitions, and Attributes Column chart.

### Taxonomy Bar Charts

(Figure 3) – In Phinch, the Taxonomy Bar Chart visualization is a highly interactive implementation of a stacked bar chart. A subway chart graphic at the top of the visualization allows users to instantaneously switch between groupings representing higher to lower level taxonomies or ontologies; the “value” and “%” buttons can be used to toggle between absolute number of observations (e.g. total number of sequence reads) or proportional abundances normalized by sample, respectively. Hovering over bars will return a pop-up box with information regarding sample metadata and taxonomy/ontology. In addition, a search box with built-in auto-completion of typed text allows users to locate a specific taxonomic or ontology term of interest within the dataset; the resulting terms can be removed from the taxonomy bar chart if needed (for example, a lab contaminant that users might want to filter out of the chart). Upon removal of a taxon or ontology term, the values or proportions displayed in the Taxonomy Bar Charts will be immediately recalculated and refreshed.

**Figure 3:**
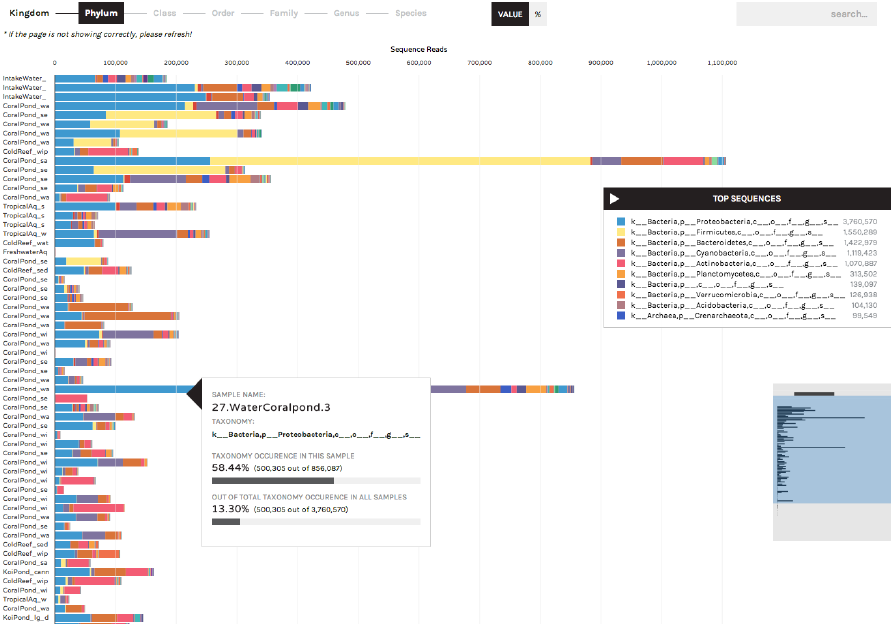
Interactive Taxonomy Bar Chart. Stacked bar charts are a common tool that researchers use to summarize data and explore biological patterns. The Phinch framework has implemented an efficient, highly interactive iteration of this research tool. Hovering over bars displays information about the sample name, observation data, and taxonomy/ontology. Users can instantly toggle between displaying the absolute number of observations (clicking the “value” button), or normalized observations (e.g. proportional abundances of genes/taxa, by clicking the “%” button). A “subway map” diagram above the bar chart enables users to summarize observation data at higher or lower levels in a taxonomy or ontology (in this example, displaying higher versus lower level taxonomy, e.g. Phylum versus Family levels). In addition, a search box with an autocomplete text feature allows users to scan the dataset for specific taxa or ontology terms.

### Bubble Chart

(Figures 4 and 5) – The Bubble Chart is a novel visualization that conveys biological information in an abstract format, aimed at facilitating objective exploration of large datasets. In this visualization, circle size is positively correlated with the abundance of a given observation within a dataset (e.g. occurrence of a taxon or ontology term). Circles can be displayed randomly in a large group (“Bubble” button), or sorted according to abundance and displayed in a linear fashion (“List” button). Clicking on a bubble allows users to explore the distribution of a taxon or ontology term across all the samples in a study (Figure 6). As in the Taxonomy Bar Charts, an auto-complete search box is also available for identifying specific terms within the data.

**Figure 4:**
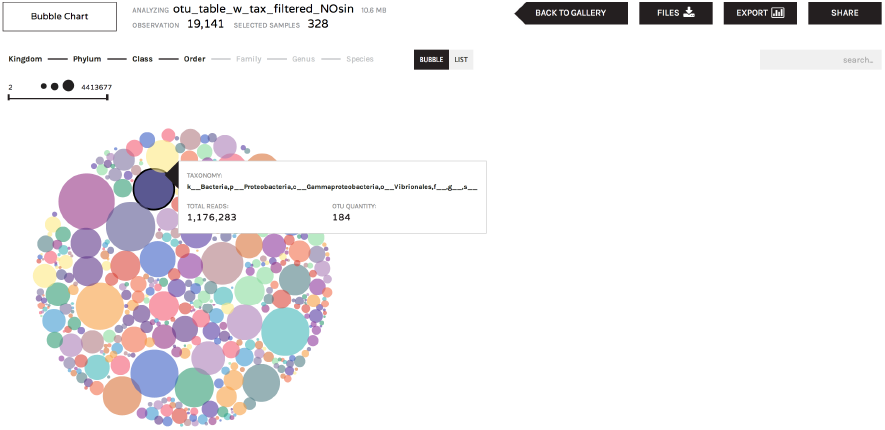
Bubble Chart Visualization highlighting Export/Share features. Bubble charts display a summary of taxonomy/ontology information, where circle size positively corresponds to the number of observations in the sample set. Hovering over each circle provides further detail. In all six visualizations, users have the ability to download and share data subsequent to visual manipulations. The “Files” button will download the filtered BIOM file along with a text log that tracks the changes and manipulations made during visual exploration. The “Export” button will download a publication-ready, high-resolution image of the onscreen visual canvas. The “Share” button is offered as a collaborative tool: it allows users to database their visualizations on the Phinch online server, giving users a unique shortlink URL which can be used anytime to directly access the saved visualization.

**Figure 5:**
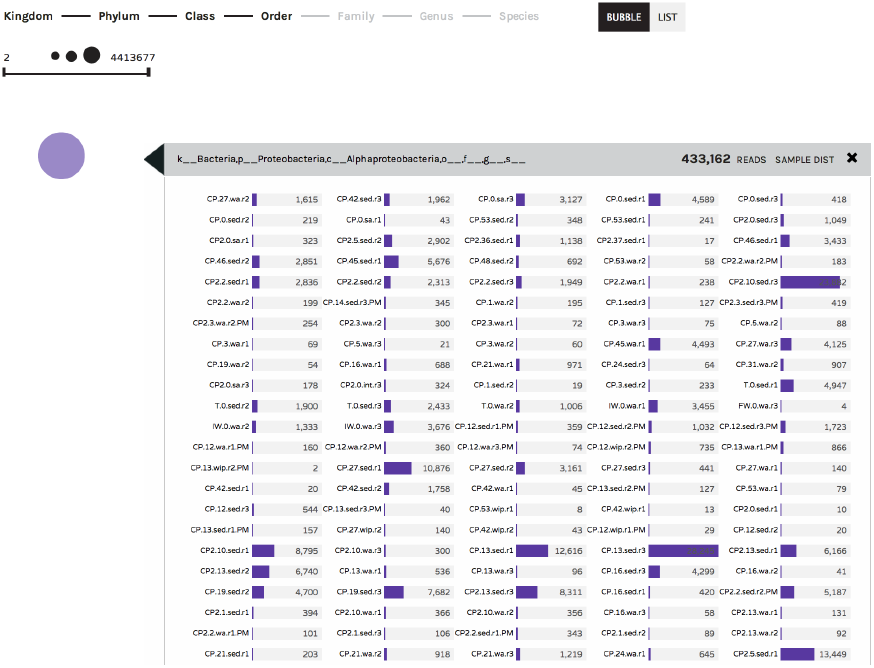
Bubble Chart interaction detail. Clicking on a colored circle within the Bubble Chart visualization will display the number of observations recorded across Sample IDs. The example below shows the number of Alphaproteobacteria 16S rRNA sequences observed across different samples in an environmental dataset.

**Figure 6:**
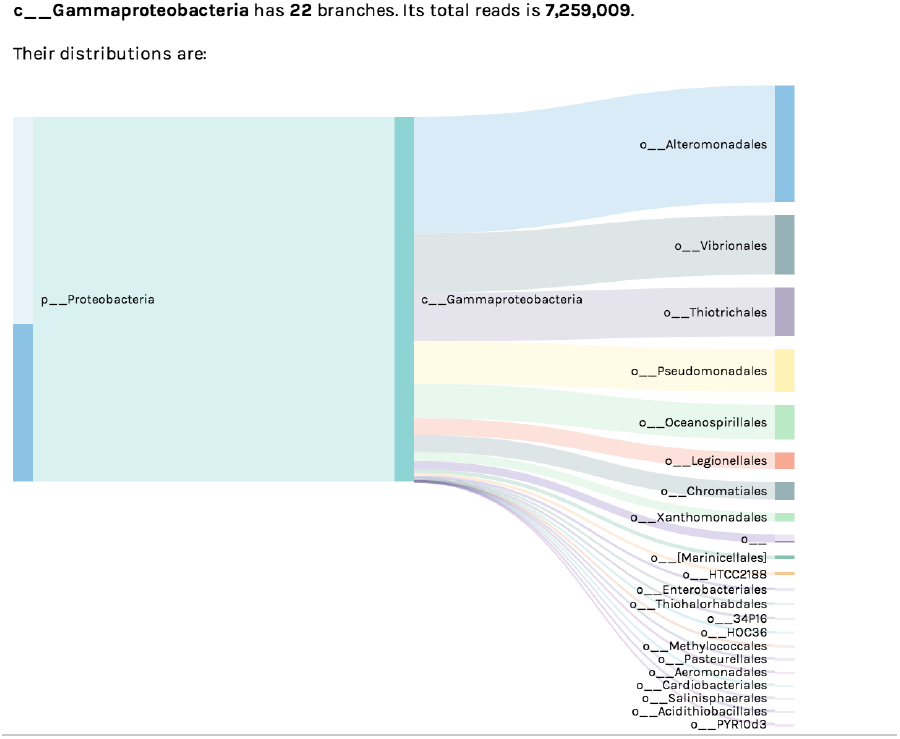
Sankey Diagram. The Sankey Diagram visualization shows the hierarchical breakdown of a dataset’s taxonomy/ontology. The pictured example shows the subset of bacterial Orders within Class Gammaproteobacteria, observed within a 16S rRNA environmental dataset. Bar width corresponds to the absolute number of sequence reads assigned to each taxon.

### Sankey Diagram

(Figure 6) – The Sankey Diagram is an organized display showing the “flow” of biological observations within a dataset, illustrating how such observations can be broken down from higher to lower level taxonomy or ontology terms. This visualization is useful for understanding what subgroups of a higher-level term are represented within a dataset (e.g. how many Orders within the Class Gammaproteobacteria have been observed, as in Figure 6), or for exploring the origin of unclassified observations (e.g. those without an assigned taxonomy or ontology term).

### Donut Partition

(Figure 7) – As currently implemented, the Donut Partition summarizes community composition (taxonomy or ontology information) for non-numerical “groupable” sample metadata. In Phinch, “groupable” metadata is classified as descriptive metadata terms used to characterize sampling sites or environmental data (e.g. water, air, soil). Clicking on a colored wedge within each donut chart will display the distribution of a taxon or ontology term across samples within that metadata category. The same or different wedges can be selected and displayed from each donut chart. The y-axis of each graph can also be set to vary across donuts (“Dynamic” button, where axes are optimized according to the observed abundances per chart), or can be set to a single standardized axis used across all donut charts (“Stand.” button).

**Figure 7:**
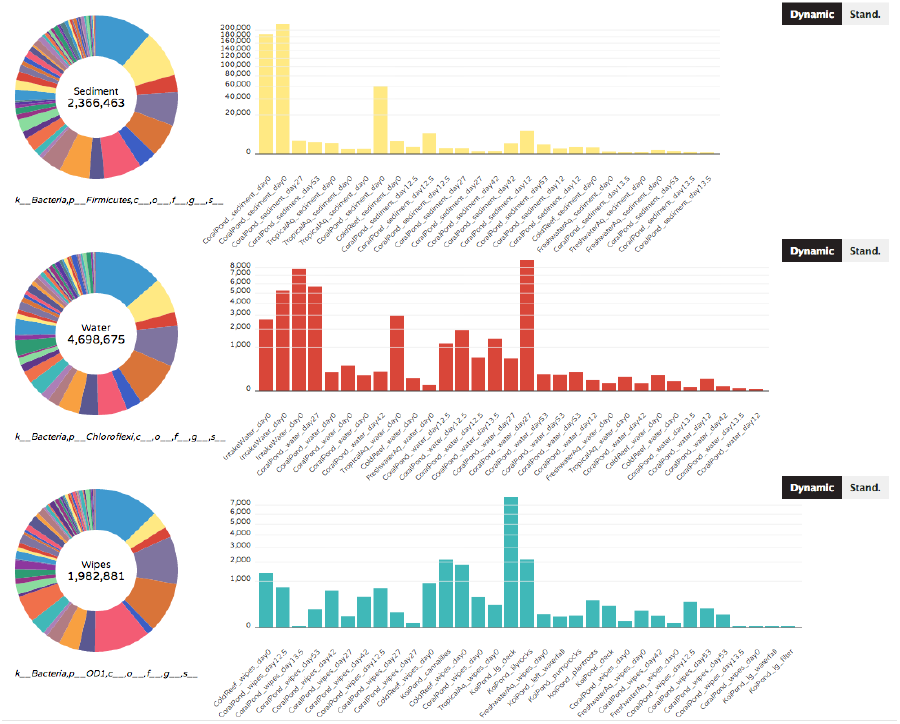
Donut Partition. The Donut Partition visualization summarizes taxonomy/ontology information according to non-numerical metadata categories (in this case, samples corresponding to different types of sample matrix). The taxonomy/ontology information can be independently displayed on each graph by clicking on the appropriate wedge of the donut chart on the left-hand side. Users can also toggle between dynamic (variable) or standardized y-axes on the displayed graphs (“Dynamic” or “Stand.” buttons, respectively).

### Attributes Column Chart

(Figure 8) - The Attributes Column Chart summarizes taxonomy or ontology terms according to numerical categories of sample metadata. Numerical categories of metadata are non-descriptive terms typically represented by environmental measurements; for example, pH readings, age, weight, temperature or other biogeochemical readings such as nitrite, ammonium, or salinity. In this visualization, the “sample distributions” box lists the sample IDs assigned to the numerical value represented by each bar. Users can immediately toggle between bar charts displaying different metadata categories, using the drop down box on the upper right hand corner of the visualization canvas (not pictured in Figure 8). As in the taxonomy bar chart, users can also toggle between absolute number of observations (“Value” button) or proportional abundances normalized by sample (“%” button).

**Figure 8:**
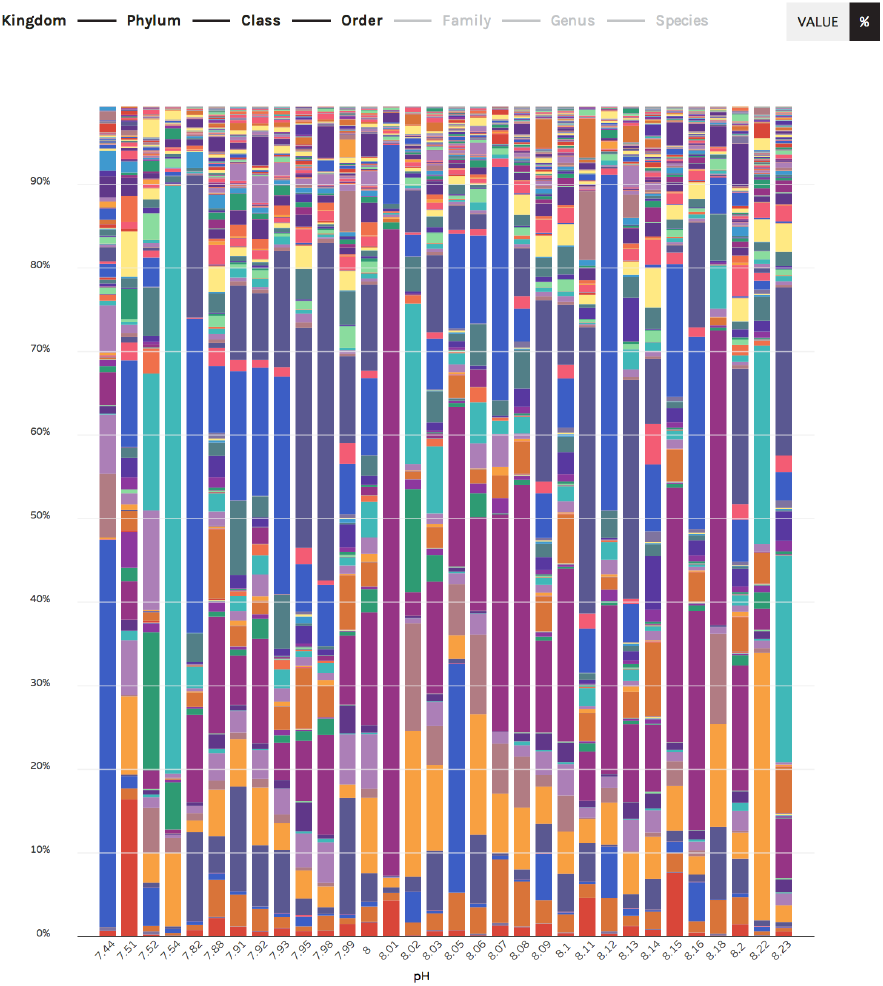
Attributes Column Chart. The Attributes Column Chart visualization summarizes taxonomy/ontology information according to numerical metadata categories (in this case, aquatic samples are grouped according to their recorded pH values). As in other visuals, the chart can be viewed at different hierarchical levels (in this case, higher versus lower level taxonomy), and users can display the stacked bars either as normalized proportional abundances (shown below) or absolute values.

After filtering data and manipulating the resulting visualizations, users have the ability to export and share their data. The “Export” button at the top of each visualization (Figure 4) allows users to download a high-resolution figure with corresponding legend, ideal for use in talks, posters or scientific publications. The adjacent “Share” button can be used to upload visualizations onto a cloud server (representing the only time datasets are transferred off a user’s local machine); the hosted data is then made available to users and collaborators via unique shortlink URL. This URL can subsequently be shared via e-mail, blog post, or even within published scientific articles, enabling direct access of data visualizations without the need to share and upload BIOM files. Finally, the “Files” button allows users to download the filtered BIOM file which underlies the pictured visualizations, as well as a text log file tracking the modifications made to the dataset during visual exploration (e.g. removing taxa from taxonomy bar charts).

## Discussion

There continues to be a persistent, vast gap between data visualization fields and the biological sciences. Despite the “growing appetite for the visual display of information… advances in visualization are not adequately described and shared with the biological community” (Wong 2012). Flexible, robust visualization capabilities such as those utilizing WebGL and HTML5 are commonly employed in the data visualization community, yet these technologies have so far seen limited applications in biological software. The Phinch framework has aimed to bridge these two areas, leveraging the critical expertise within each field via an interdisciplinary team of visual artists, software engineers, and practicing biologists. The development process has aimed to incorporate the existing body of knowledge related to effective software design (human-computer interaction, color theory, visual display of information – see http://www.infovis-wiki.net/index.php?title=Category:Publications) in pursuit of a long-term, scalable framework for scientific visualizations of –Omic datasets.

Phinch is unique, in that it represents a scientific visualization framework written directly for a web browser. Developing software within such a framework has only been made possible in recent years due to significant advances in web browser capabilities. This is in stark contrast to past scenarios, where similar visualization software would need to be developed as a downloadable standalone application, with a limited user base restricted to a specific operating system(s). By targeting a web browser instead, Phinch aims to bring data visualization tools to a much larger audience of users; the software is agnostic to operating system and can be readily utilized by anyone with access to a modern web browser.

While developing Phinch, we have aimed to minimize overlap with existing data analysis pipelines. The implemented visualization tools were designed to improve on current software, and provide additional functionality and features that facilitate rapid exploration and identification of biological patterns within large –Omic datasets. The Phinch data visualization framework represents a significant advance over alternative tools, and provides proof-of-concept for the use of data visualization as a routine part of the scientific data analysis workflow. Thus, the novelty of Phinch lies in the fact that it emphasizes interactivity (unlike other biological software tools that largely generate static visualizations), provides an exploratory framework that leverages visual displays of abstract data (emphasizing human cognition and quick, in-depth exploration of large data volumes), promotes accessibility to diverse audiences of researchers and the public alike (users do not need a deep level of computational knowledge or programming skills to access data visualizations), and facilitates both collaboration and efficient science (via implemented sharing and export features).

## Supplementary File 1: Preparing BIOM files for visualization

Phinch currently supports downstream analyses of BIOM files, Biological Observation Matrix format (BIOM version 1.0, http://biom-format.org/), a JSON file type (denoted using the .biom file extension) used to represent diverse types of genomic data. The most typical user applications are environmental rRNA amplicons or shotgun metagenomic data, although any type of sample/observation data can be represented as BIOM files (RNA-seq, gene variants, morphological character matrices, etc.)

To prepare files for visualization with Phinch, follow these steps:

### Step 1: Prepare a QIIME-style mapping file for sample metadata

Sample metadata is defined as any descriptive information about a biological sample or the environment where the sample was collected; any type of metadata can be included that may be useful for interpreting and analyzing patterns in a dataset. Some common types of sample metadata include geographic coordinates (latitude/longitude), collection date, state/country, sampling matrix (water, air, soil, sediment), treatment group, etc. Mapping files can contain as much or as little sample metadata as is useful or necessary. For example, sample metadata for a human microbiome study might also include information about patient gender, body site where samples were collected, or patient age. Mapping files should be prepared according to these QIIME guidelines: http://qiime.org/documentation/file_formats.html

**To label samples in Phinch and export graphics with human-readable IDs, include a column in the metadata mapping file with a header named “phinchID”** (these entries can be the same or different as the first SampleID column). The phinchID values will be pulled through into the visualizations to populate graph axes. If this column is not included, an arbitrary numerical ID will be assigned to each sample. **For optimal visualization, phinchID strings should be no longer than 15 characters**.

For columns containing numerical metadata values: Enter **‘no_data’** if there is no measurement available for a given sample (for example, columns listing temperature, pH, or other chemical measurements where a given value was not recorded for some samples). Blank fields within a column are not recommended, since this may lead to improper importing and formatting of metadata values.

For an example sample mapping file, refer to https://github.com/PitchInteractiveInc/Phinch/wiki/Quick-Start

#### Some notes on metadata formatting

In order to be properly detected, all date/time metadata must adhere to the MIxS standardized format (http://wiki.gensc.org/index.php?title=MIxS) and entered into one column in the original sample metadata mapping file, using the following convention:

#### [YYYY]-[MM]-[DD]T[hh]:[mm]:[ss]-[Z]

This date format lists the year, month, and day, followed by a 24hr timestamp with a UTC offset (Z). Inclusion of timestamp and UTC offset are both optional; metadata columns can include date only. For example, metadata for a sample collected at 2:30pm EST on May 4, 2007 would be entered as: 2007-04-05T14:30:00-05:00 Similarly, any geographic coordinates or GPS data must be entered as decimal degrees (the format used by GoogleMaps, e.g. -90.017926). We recommend using separate columns labeled “Latitude” and “Longitude” in your original sample metadata mapping file, to ensure that GPS metadata is correctly detected.

For numerical values with units, there should be a space inserted between each value and unit. For example, altitude data should be entered as 2421 m, instead of 2421 or 2421m.

### Step 2: Prepare biological matrix data as a BIOM file

BIOM files are now the default output for rRNA amplicon workflows (OTU picking - http://qiime.org/tutorials/otu_picking.html) and analysis of shotgun metagenome data (http://qiime.org/tutorials/shotgun_analysis.html) in the QIIME software package (http://qiime.org).

In QIIME (version 1.7 or later), users can prepare a BIOM file for visualization by executing the following commands.

#### First, construct an OTU table

~~~
make_otu_table.py -i final_otu_map_mc2.txt -o otu_table_mc2_w_tax.biom -t
rep_set_tax_assignments.txt
~~~

Where the input file (-i) is an OTU Map (defining clusters of raw sequence reads), and the taxonomy file (-t) contains the taxonomy or ontology terms that correspond to each observation.

#### Second, add sample metadata to the BIOM file

**All sample metadata and taxonomy/ontology information MUST be embedded in the BIOM file before being loaded into Phinch.**

In QIIME version 1.8 this can be done using the following command:

~~~
biom add-metadata -i otu_table_mc2_w_tax.biom -o
otu_table_mc2_w_tax_and_metadata.biom -m sample_metadata_mapping_file.txt
~~~

In QIIME version 1.7 or below, metadata can be added using the following command:

~~~
add_metadata.py -i otu_table_mc2_w_tax.biom -o
otu_table_mc2_w_tax_and_metadata.biom -m sample_metadata_mapping_file.txt
~~~

The input file (-i) is the BIOM file from the previous step, and the mapping file (-m) is the tab-delimited mapping file prepared in Step 1 (formatted according to QIIME instructions: http://qiime.org/documentation/file_formats.html)

### Step 3: Upload BIOM file with embedded observation/sample data at http://phinch.org, using the Google Chrome browser

#### File Conversion Instructions: tab-delimited or matrix data to BIOM format

To visualize biological data currently formatted as a tab-delimited text file (e.g. the style of OTU tables produced by older versions of QIIME, or any other type of genomic/morphological data that can be represented in matrix format, refer to the BIOM documentation for conversion instructions (http://biom-format.org/documentation/biom_conversion.html). Phinch supports both “sparse” and “dense” BIOM formats (although sparse BIOM files are highly recommended, since the file size is much smaller). Full documentation for the BIOM file format can be found at http://biom-format.org

